# An interactive web application for exploring human plasma and fibroblast metabolomics data from patients with inborn errors of metabolism

**DOI:** 10.1101/2023.12.11.571124

**Authors:** Ling Cai, Hieu S Vu, Wen Gu, Hongli Chen, Jordan Franklin, Lea Abou Haidar, Zheng Wu, Chunxiao Pan, Feng Cai, Phong Nguyen, Bookyung Ko, Chendong Yang, Lauren G. Zacharias, Jessica Sudderth, Sarah Montgomery, Crescenda Uhles, Heather Fisher, Julianna Hudnall, Callie Hornbuckle, Christine Quinn, Donnice Michel, Luis Umaña, Angela Scheuerle, Markey C. McNutt, Garrett K. Gotway, Bushra Afroze, Min Ni, Ralph DeBerardinis

**Author notes:** These authors contributed equally (Ling Cai, Hieu S Vu).

## Abstract

Metabolomic profiling is instrumental in understanding the systemic and cellular impact of inborn errors of metabolism (IEMs), monogenic disorders caused by pathogenic genomic variants in genes involved in metabolism. This study encompasses metabolomics analysis of plasma from 474 individuals and fibroblasts from 67 subjects, incorporating healthy controls, patients with 65 different monogenic diseases, and numerous undiagnosed cases. We introduce a web application designed for the in-depth exploration of this extensive metabolomics database. The application offers a user-friendly interface for data review, download, and detailed analysis of metabolic deviations linked to IEMs at the level of individual patients or groups of patients with the same diagnosis. It also provides interactive tools for investigating metabolic relationships and offers comparative analyses of plasma and fibroblast profiles. This tool emphasizes the metabolic interplay within and across biological matrices, enriching our understanding of metabolic regulation in health and disease. As a resource, the application provides broad utility in research, offering novel insights into metabolic pathways and their alterations in various disorders.

## Introduction

Metabolomic profiling of plasma represents a foundational tool in the study of human metabolism. It serves as a gateway to decipher the genetic underpinnings of metabolic traits and advance precision medicine within both healthy and disease-affected populations. Numerous studies have leveraged this approach to enhance biomarker discovery, refine diagnostic protocols, and deepen our understanding of pathophysiology [1-6].

Fibroblast cultures derived from patient skin biopsies also provide useful resources in genetic disease diagnosis and research. Their ease of acquisition and the ability to harmonize the culture environment to identify cell-autonomous metabolic properties make fibroblasts a helpful complement to plasma. The metabolic differences among fibroblasts derived from different individuals arise from intrinsic cellular processes rather than the effects of diet, medications and other host factors reflected in the plasma metabolome. Despite their value, large-scale metabolic profiling of fibroblast lines remains underexplored.

In this study, we employed metabolomics profiling to analyze the plasma of 474 individuals and fibroblast cultures from 67 subjects recruited through the Genetic and Metabolic Disease Program (GMDP) at UT Southwestern Medical Center, including cohorts from a tertiary-care pediatric hospital in Dallas, Texas, a biochemical genetics clinic at the Aga Khan University Hospital in Karachi, Pakistan, and collaborators from other centers. This undertaking involves a concurrent examination of plasma and fibroblast metabolomes in a human cohort highly enriched for IEMs. While a companion paper details scientific insights derived from these analyses[7], herein we present an interactive web application (https://lccl.shinyapps.io/GMDP/) designed to enable user-driven exploration and analysis of these complex datasets.

## Materials and Methods

### Human Subjects

This database compiles data derived from research that involved human subjects. The detailed methodology, including participant recruitment, data collection procedures, and ethical considerations, are comprehensively described in a companion paper[7]. The clinical research protocol STU112014-001 received full approval from the University of Texas Southwestern Medical Center Institutional Review Board and the study is registered on ClinicalTrials.gov as NCT02650622.

### Metabolomics data acquisition and processing

The plasma metabolomics data were acquired in two batches whereas the fibroblast data were acquired in a single run on a high-resolution mass spectrometry (HRMS) - 6550 iFunnel Q-TOF mass spectrometer (MS) (Agilent Technologies, CA) coupled with a 1290 UHPLC system for reverse-phase chromatography. The MS was operated in both positive and negative (ESI+ and ESI-) modes.

Metabolites that were not detected in ten or more samples were removed. In instances where metabolites were identified in both ionization modes or had multiple ions representing the same metabolite, we retained only the ion with the highest signal intensity. Quality control (QC) samples, which consisted of a run-specific mixture of aliquots from all samples and a historical QC sample used across multiple runs, were injected at regular intervals interspersed among the experimental samples. These QC samples facilitated a robust LOESS (locally estimated scatterplot smoothing) signal correction, normalizing the sample signals against a QC-derived trendline to account for any temporal drifts [8]. Following this correction, we established a normalization factor from a subset of stable metabolites—those with low variation, moderate to high signal intensity, and a signal-to-noise ratio exceeding five—to adjust the signal based on the ‘total stable ion count’. Subsequently, the data underwent a log_2_ transformation. To integrate the two batches of plasma data, we aligned them by their common metabolites and applied the ComBat method from the ‘sva’ R package [9] for batch correction, confirmed by a validation set from overlapping samples in both batches.

### Web application construction

Our interactive web application was built by R shiny, with the following R packages: “data.table”[10], “dplyr”[11], “openxlsx”[12], “reshape2”[13], “stringr”[14], “tidyverse”[15] for data wrangling and preparation; “Hmisc”[16], and “nlme”[17] for statistics; “ape”[18], “ComplexHeatmap”[19], “InteractiveComplexHeatmap” [19], “ggplot2”[20], “ggpubr”[21], “ggrepel”[22], “patchwork”[23], “pathview”[24], “plotly”[25], and “RcolorBrewer”[26] for visualization; “bslib”[27], “DT”[28], “htmltools”[29], “shinycssloaders”[30], “shinyjs”[31], and “shinyWidgets”[32] for app user interface and server construction. The app is deployed to https://www.shinyapps.io/ through “rsconnect”[33]. The processed data are available for download from the web application. Code for the web application is available at the GitHub repository https://github.com/cailing20/GMDP_app.

### Statistics

All analyses were performed with R version 4.0.2 [34]. For controlling the false discovery rate (FDR) in multiple testing scenarios, we applied the Benjamini-Hochberg procedure. We utilized a robust Z-score formula, given by 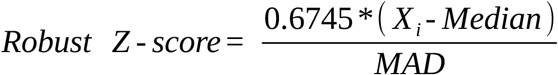, where MAD = median (|X-median|). This approach differs from the traditional Z-score that is based on the mean and standard deviation, providing a measure less sensitive to outliers.

Pearson correlation is used by default, with Spearman rank correlation available as an alternative in the correlation-based tools. The examples presented here use Pearson correlation unless otherwise specified.

For the analysis comparing diagnosed and reference samples, we employed a linear mixed-effects model using the “nlme” package in R [17]. This model incorporates variability between replicates as random effects, while disease status is treated as a fixed effect, allowing for a nuanced assessment of the data.

## Results

We organized the 13 analysis tools in the web application into four domains – “data overview”, “metabolic deviation”, “metabolic neighbors”, and “plasma-vs.-fibroblast comparison” (Figure 1). The tutorial section of the app provides snapshots with notes on how to use each utility.

**Figure 1.**
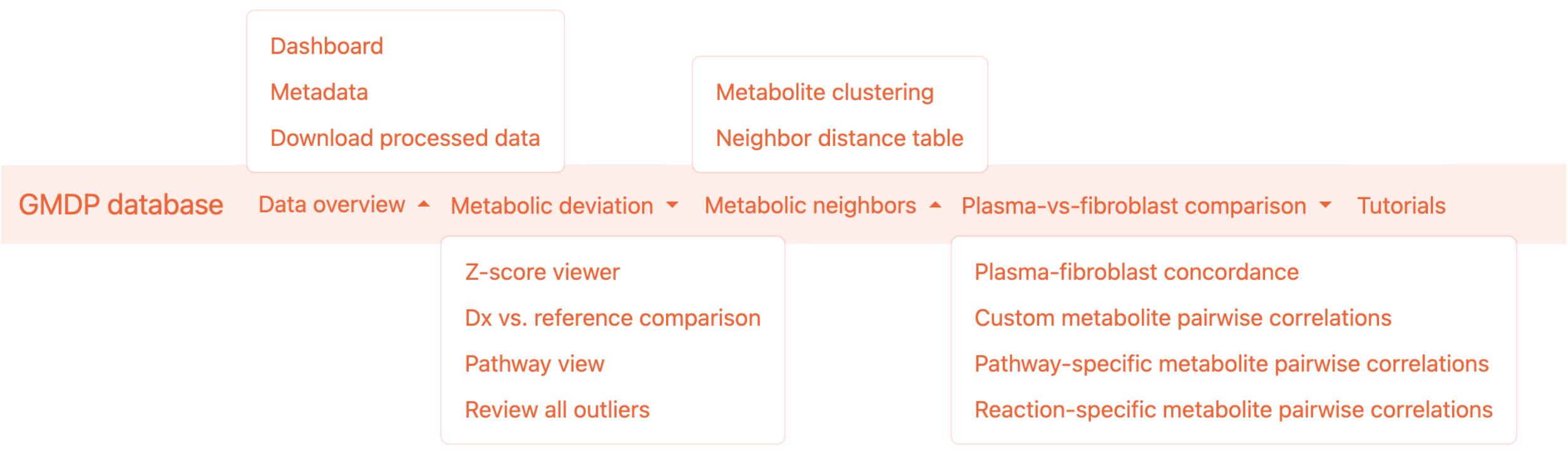
Layout of Tools in the GMDP Database App. This figure displays how the 13 tools in the GMDP app are grouped into four main sections. The tutorial section provides example usage of each functionality.

### Data Overview

#### Dashboard

The “Dashboard” tab presents an overview of the plasma and fibroblast datasets, categorized by diagnosis status (Figure 2). It also aggregates additional demographic details for the plasma dataset, including pedigree, age, ethnicity, and race. In the plasma dataset, 59% of the participants are healthy, while the fibroblast dataset includes 13% healthy individuals. The inclusion of these reference samples is crucial as they provide a baseline for identifying metabolic deviations associated with diseases.

**Figure 2.**
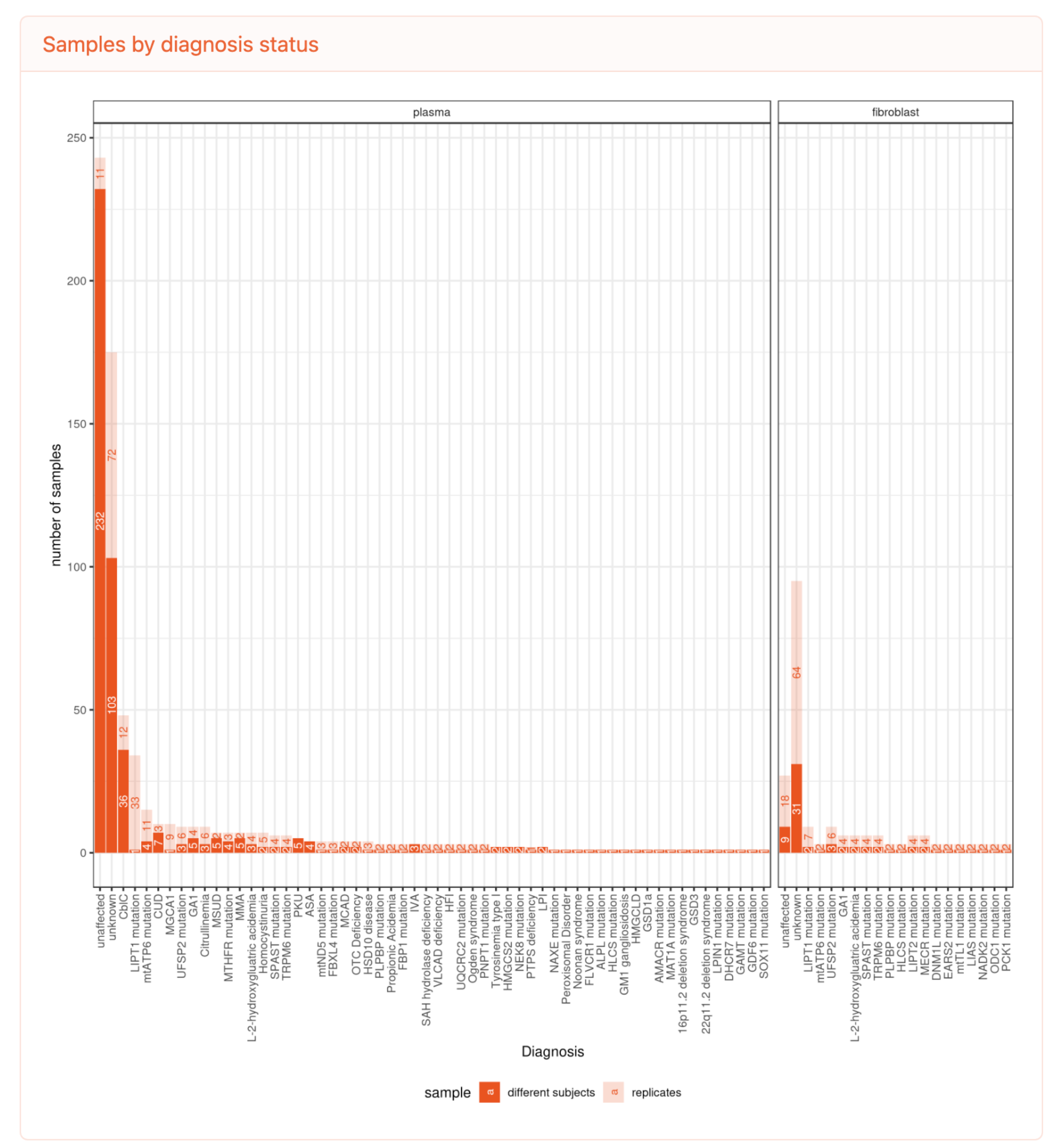
Plasma and Fibroblast Sample Distribution by Diagnosis Status in Data Summary Page. The chart illustrates the count of samples by diagnosis status for plasma and fibroblast datasets. The plasma dataset includes samples from 232 unaffected subjects, 103 subjects with undiagnosed diseases, and 139 subjects with 56 different genetic diseases. The fibroblast dataset comprises samples from 9 unaffected subjects, 31 subjects with undiagnosed diseases, and 27 subjects with 18 different genetic diseases. The plasma replicates encompass technical replicates and samples taken at different times from the same subject. Most fibroblast samples were processed in technical triplicates.

#### Metadata

In the “Metadata” tab, users can access detailed sample and metabolite information from both plasma and fibroblasts. This section includes hyperlinks to external databases, offering resources for further exploration into disease contexts and metabolite profiles (refer to Figure S1). For comprehensive analysis, the fully processed datasets for plasma and fibroblasts are available for download in the “Download processed data” tab.

### Metabolic Deviation

Metabolic variability arises from many factors, including fasting status, age, sex, race, environment, and lifestyle. IEMs produce larger and more consistent deviations from the range observed in healthy subjects, and these large deviations are the primary focus of our analysis.

#### Z-score Viewer

The “Z-score Viewer” tool provides a robust analysis of individual sample variations by utilizing Robust Z-scores, which are less sensitive to data skewness due to their reliance on the median and median absolute deviation, rather than the mean and standard deviation. This adjustment is particularly vital in our context, where disease samples with very large metabolite deviations from the healthy range would skew the mean and standard deviation. This tool allows for a detailed exploration of these scores for both individual metabolites and samples. An application of this is demonstrated in Figure 3, where outliers in phenylalanine levels are identified (left), highlighting subjects with phenylketonuria (PKU) due to mutations in the phenylalanine hydroxylase (PAH) enzyme. On the right, all detected metabolites from a patient with PKU are ranked by robust Z-score.

**Figure 3.**
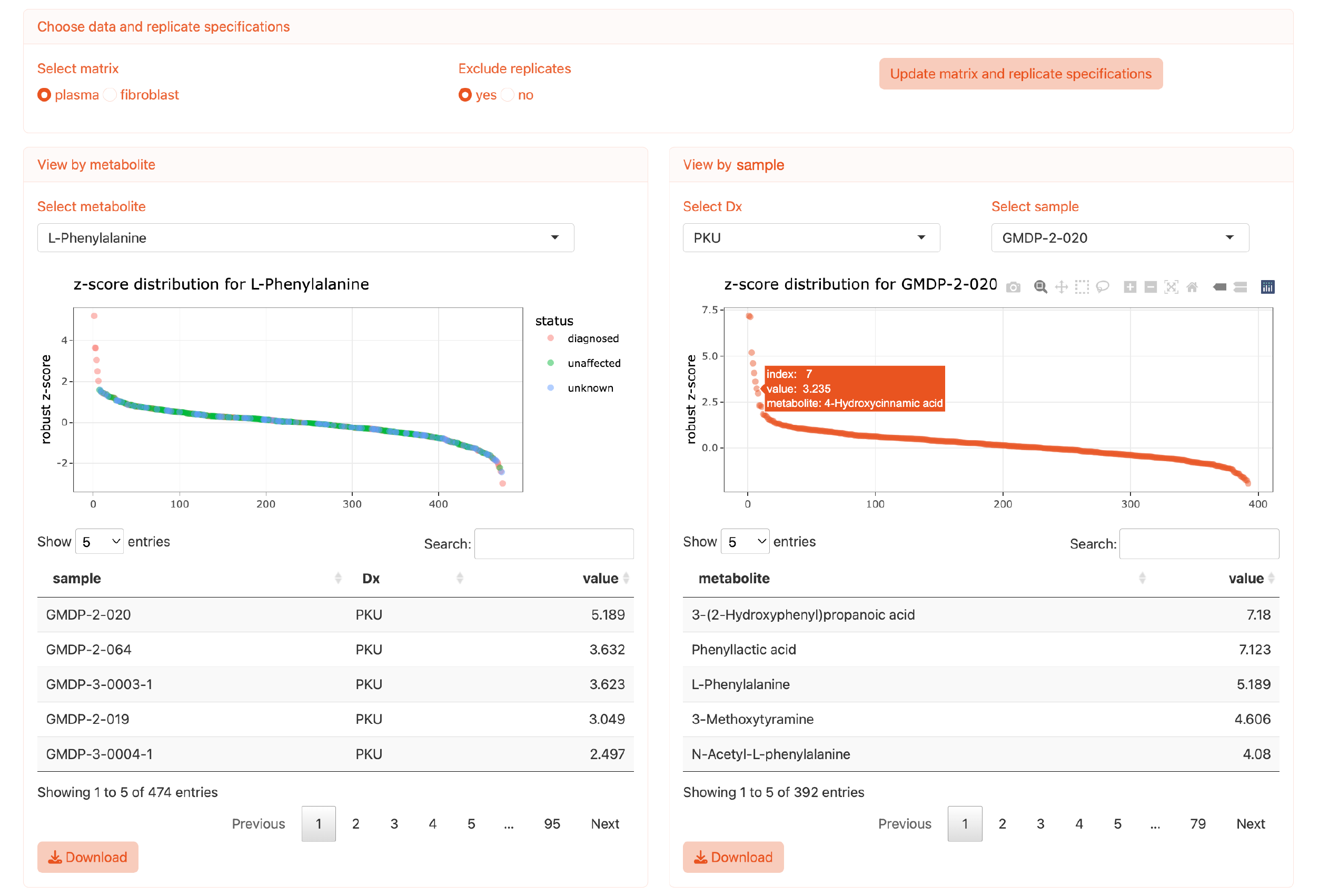
Interactive Z-Score Viewer for Metabolite Analysis. This interactive tool allows users to visualize robust z-scores for specified metabolites across samples (left panel) or for individual samples across metabolites (right panel). Users can tailor the analysis by selecting the data matrix and specifying whether to include replicates. The dot plots are interactive, providing detailed insight upon selection. Data from the analysis can be downloaded directly from this interface.

#### Dx vs. Reference Comparison

The “Dx vs. Reference Comparison” tool facilitates group analysis, comparing disease samples with reference groups to ascertain metabolic discrepancies. This function accommodates the simultaneous comparison of multiple diseases, which can be particularly helpful when diseases share similar metabolic profiles. Diseases are visually differentiated in the dot plot by color coding, enhancing ease of interpretation. For instance, as depicted in Figure 4a, fibroblast samples from subjects with three lipoylation defects—*LIPT1, LIPT2*, and *LIAS* mutations—demonstrate a consistent decrease in isovalerylcarnitine compared to the “unaffected and other diagnosed” reference group, indicating a metabolite anomaly that unifies these conditions.

**Figure 4.**
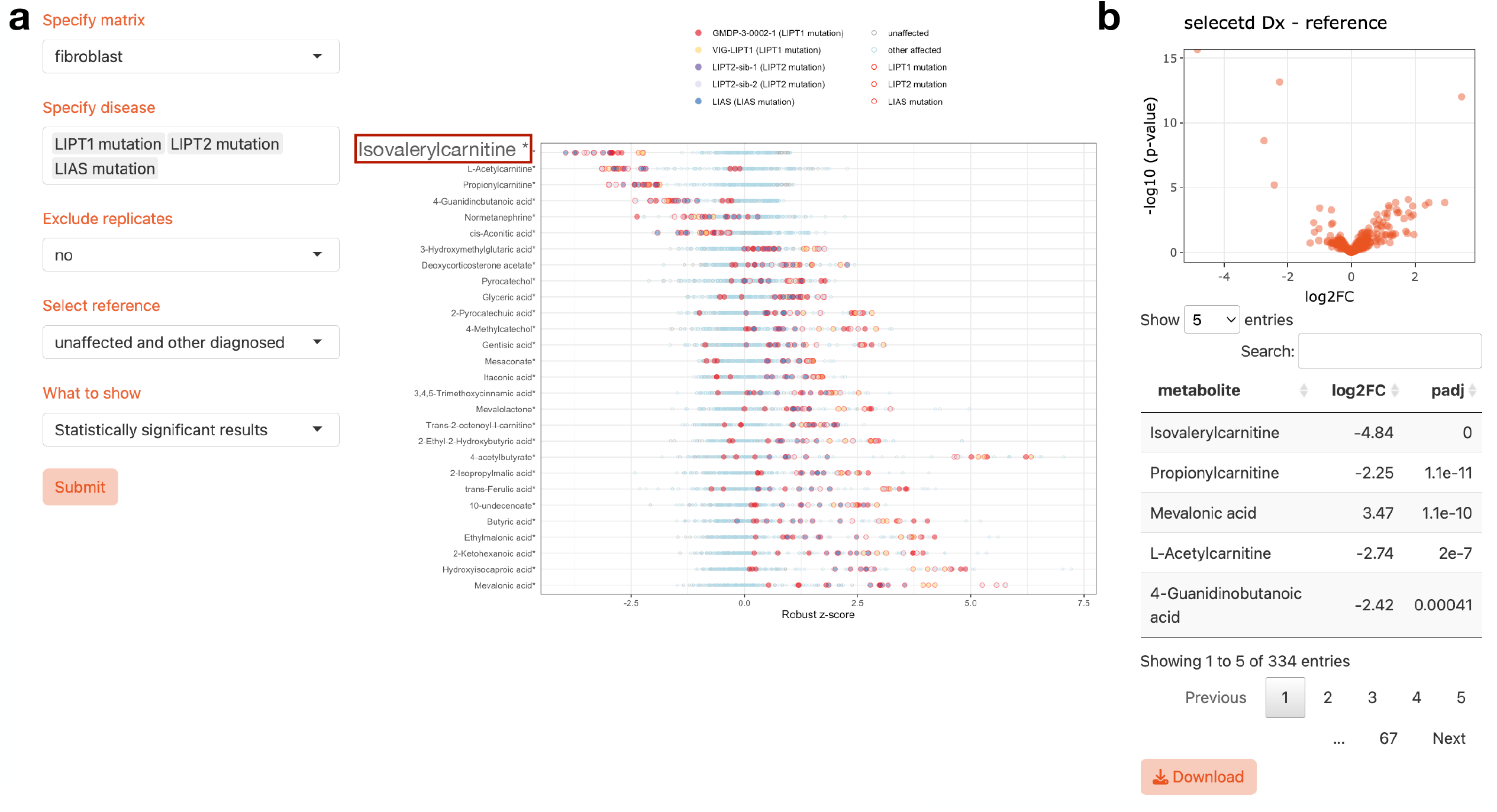
Usage example of the Dx vs. reference comparison. **a**, This figure demonstrates a robust z-score dot plot and statistical comparison between subjects with various diseases and reference samples. Users can select a single disease or multiple diseases simultaneously for comparison against reference groups, which can be exclusively unaffected samples or a combination of unaffected and other diagnosed conditions. The color-coding in the dot plot differentiates between the subjects’ diagnoses. There is flexibility to display or exclude replicates in the plot. All data, inclusive of replicates, is used in the statistical analysis with untransformed values. Metabolites with significant variance from the reference, identified post-adjustment for multiple comparisons via the Benjamini-Hochberg method, are highlighted with an asterisk. The underlying statistical approach employs a linear mixed-effects model, which treats disease status as a fixed effect while incorporating cross-replicate variability as random effects. **b**, For the volcano plot, p-values below the precision limit are capped at 2.2e-16, although they are documented as zero in the statistical summary table.

Users have the option to customize the display of data points in the dot plot, such as deciding whether to include multiple replicates or time points for a given subject. This feature allows the user to assess variability in cases where multiple samples are available from the same patient (e.g. the LIPT1-deficient patient in Figure 4a). Alternatively, for diseases with larger sample sizes, like the Cobalamin disorder CblC with 36 subjects, users can opt to streamline the visual presentation by excluding replicates.

Underlying these visual representations, the robust Z-score is utilized for individual data point placement, while group comparison statistics are derived from untransformed data through a linear mixed-effect model. This model accounts for intra-sample variability as random effects and the differences between the disease and reference groups as fixed effects. The outcome of this statistical treatment feeds into both the volcano plot and a downloadable table, providing users with a comprehensive data interpretation, as seen in Figure 4b.

For diseases with a smaller sample size, where few metabolites meet the stringent threshold for statistical significance (adjusted p-value < 0.05), an additional display option is available. Users can select to view the “Top & bottom 25” metabolites, facilitating a focused review of those with the most pronounced deviations from the median of the reference group. This option offers a targeted approach to discern potential biomarkers or metabolic disruptions, as illustrated in Figure S2.

#### Pathway View

The “Pathway View” panel of the “Metabolic Deviation” module offers a focused examination of metabolic pathways, displaying either log_2_ fold change (log_2_FC) from group comparisons or robust Z-scores from individual assessments. This tool allows users to visualize how metabolites differ between diseased and healthy states within specific metabolic pathways. Figure 5 showcases the “Pathway View” function, highlighting metabolic discrepancies within the valine, leucine, and isoleucine degradation pathway when comparing subjects with maple syrup urine disease (MSUD) to healthy individuals. Accumulation of intermediates preceding the enzymatic reaction mediated by the Branched Chain Keto Acid Dehydrogenase E1 Subunit Alpha (BCKDHA) aligns with the known metabolic bottleneck characteristic of MSUD. This tool can also reveal metabolic variances on an individual level (Figure S3), where a single subject with MSUD is analyzed, demonstrating a similar pattern of metabolic disruption.

**Figure 5.**
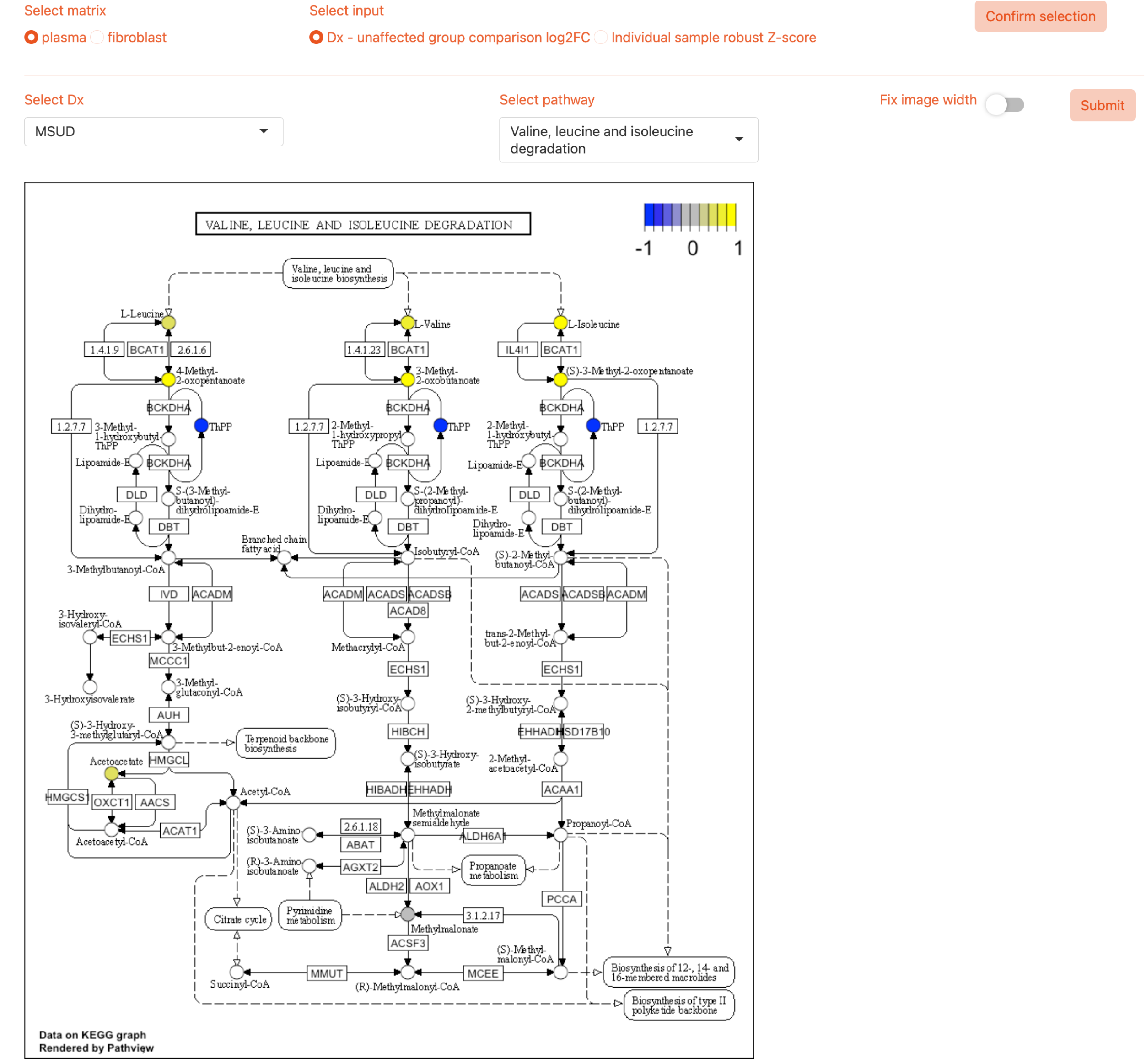
Examination of MSUD metabolic deviation in the branched-chain amino acid catabolic pathway. This figure depicts a targeted pathway analysis for Maple Syrup Urine Disease (MSUD), focusing on the degradation of the branched-chain amino acids (BCAAs) valine, leucine, and isoleucine, which accumulate in MSUD due to defective catabolic enzymes. The data presented are derived from the plasma samples group comparison, contrasting MSUD subjects against unaffected controls. Metabolites covered by our panel within the pathway are annotated with a color-coded indicator based on the log2 fold change (log2FC) values, which reflect the relative abundance in the MSUD plasma samples.

For a more expansive analysis that does not require a predefined hypothesis, the “Metabolic Pathway Panorama” option provides a comprehensive metabolic landscape. This broad view, illustrated in Figure S4, encompasses the full spectrum of metabolite variations within the context of a particular disease or individual sample.

#### Review All Outliers

The “Review All Outliers” function in the “Metabolic Deviation” module enables users to identify and analyze metabolic outliers with robust Z-scores exceeding a threshold of absolute value greater than 3. This tab provides an inclusive view, allowing for targeted filtering by disease, metabolite, or subject ID, effectively homing in on relevant data points. For instance, analysis within this tool revealed elevated 4-acetylbutyrate in fibroblast lines from patients with a subset of mitochondrial disorders, as shown in Figure S5a, and consistently high methylmalonic acid levels in plasma from subjects with methylmalonic acidemia or cobalamin processing defects, as seen in Figure S5b. This functionality simplifies the process of detecting and investigating metabolic anomalies across diverse datasets.

### Metabolic Neighbors

#### Metabolite Clustering

Understanding the complex interactions between metabolites in biological systems can provide valuable insights into metabolic pathways and their regulation. The “Metabolite Clustering” feature allows users to delve into these interactions by examining correlations among metabolites in plasma and fibroblast datasets. By setting a custom threshold for correlation strength, users can visualize how metabolites group together in a circular dendrogram format (Figure 6). This visual representation enables the identification of subclusters of metabolites that share similar properties or functions within the plasma data.

**Figure 6.**
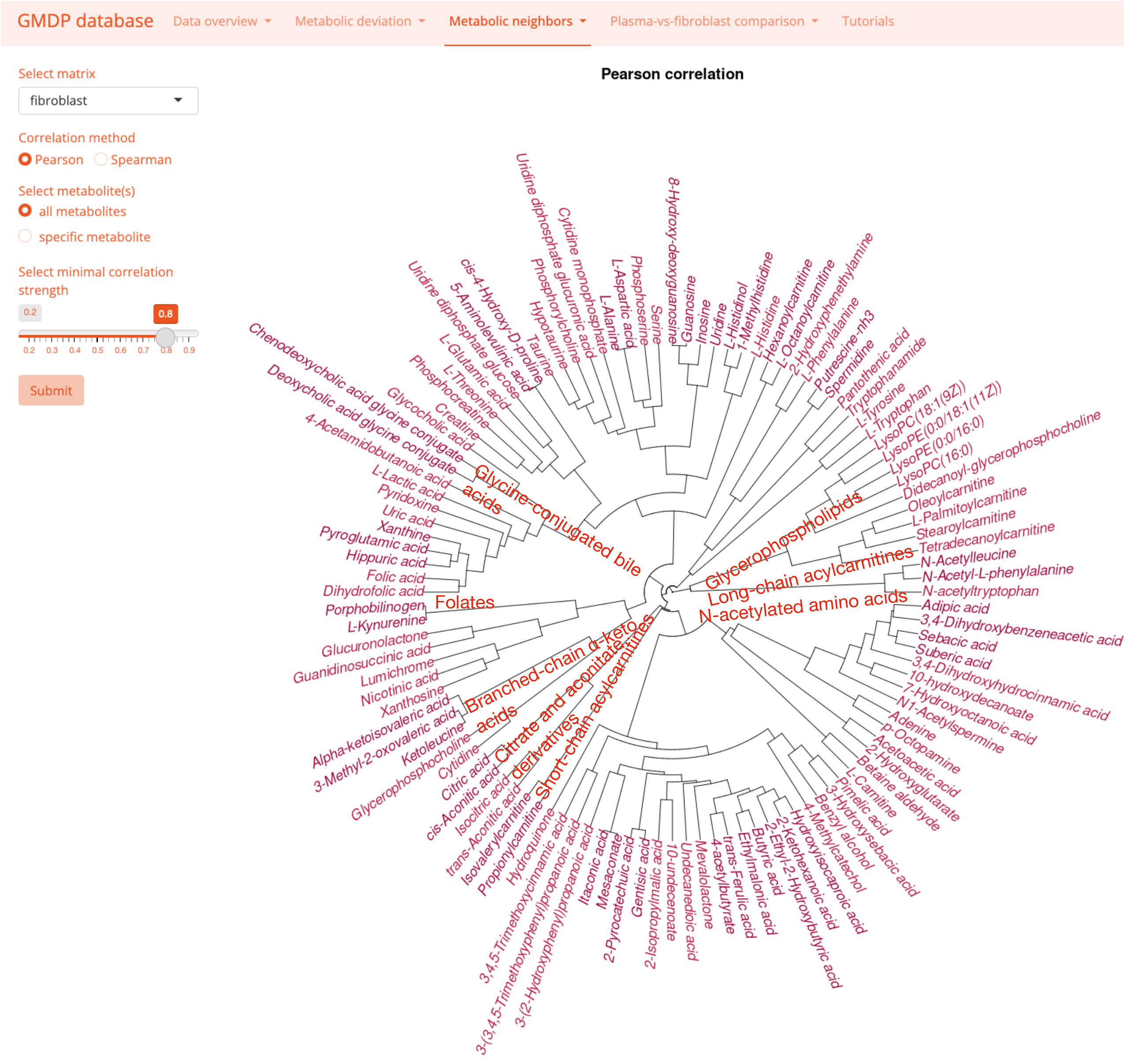
Clustering of metabolites in fibroblast dataset. Circular dendrogram illustrating the hierarchical clustering of metabolites measured in fibroblasts. The dendrogram is constructed using average linkage hierarchical clustering based on the absolute correlation distance between metabolites, with a minimum correlation strength threshold as defined by the user. Notable clusters were annotated to indicate groups of metabolites with related biochemical properties or shared metabolic pathways. This visualization helps in identifying potentially related metabolites based on their co-expression patterns, which can provide insights into metabolic relationships and network dynamics within the plasma metabolome. The dendrogram’s threshold for correlation strength can be adjusted, affecting the clustering granularity. Metabolites below the threshold are not visualized to maintain focus on stronger and potentially more biologically relevant correlations.

However, when a lower threshold for correlation strength is chosen, the resulting dendrogram may become saturated with connections, making it difficult to discern individual relationships. To address this, we offer an additional functionality that allows users to focus on a specific metabolite of interest. This generates a clustered heatmap, displaying a subset of metabolites that closely associate with the selected one. Figures S6 and S7 demonstrate this application, revealing how isovalerylcarnitine clusters with other short-chain carnitines. This pattern is consistent across both fibroblast and plasma samples, with disease samples often presenting as outliers.

#### Neighbor Distance Table

Complementing the “Metabolite Clustering” is the “Neighbor Distance Table”, which provides a tabular approach to analyze metabolic similarities. This feature is designed for users who prefer a more quantitative analysis over the visual dendrograms. It utilizes two distance metrics—Euclidean and absolute correlation distance—to measure the closeness between pairs of samples or pairs of metabolites. Users can tailor the analysis to specific datasets and samples, facilitating a bespoke exploration of the data.

For instance, when investigating the metabolic neighbors of propionylcarnitine within plasma samples, the table reveals that methylmalonic acid—a key intermediate in propionate metabolism—surfaces at the top of the list, followed by a series of short-chain acylcarnitines (Figure S8a). This suggests a shared metabolic pathway or a common regulatory mechanism. Similarly, when identifying fibroblast samples that resemble those with the LIPT1 mutation, the table lists top matches from other subjects with defects in lipoylation and other mitochondrial activities, indicating a shared disease phenotype despite the genetic diversity (Figure S8b).

### Plasma-vs-fibroblast comparison

#### Plasma-Fibroblast Concordance

In the ‘Plasma-fibroblast concordance’ analysis depicted in Figure S9, concordance between plasma and fibroblast metabolomes is quantitatively assessed through Pearson correlations at two levels: across shared metabolites and shared subjects. Our dataset includes 42 subjects with shared profiles, which comprises 27 undiagnosed individuals, one healthy subject, and 14 diagnosed subjects across eight diseases. The tool’s correlation analysis reveals that statistically significant concordance between plasma and fibroblast samples is rare, with few metabolites showing significant correlation in shared subjects, and conversely, no individual subjects displaying significant correlation across shared metabolites. The correlation outcomes are often skewed by outliers, particularly in disease-affected samples, suggesting distinct underlying metabolic mechanisms between plasma and fibroblast samples. This observation is critical, indicating that each matrix may reflect different aspects of the metabolic phenotype, especially in pathological conditions, thus challenging the assumption of direct comparability of plasma and fibroblast metabolomic data.

#### Custom Metabolite Pairwise Correlations

The correlation between a pair of metabolites can provide insights into potential metabolic pathways or biological processes that are shared between the two metabolites. Surprisingly, the concordance between plasma and fibroblast data over 44,850 pairwise correlations for 300 shared metabolites is rather weak (Figure 7a), suggesting the factors underlying the metabolic covariations (such as metabolic connectivity) differ between the two systems. We also observed that the correlations in fibroblast samples are more robust than from the plasma dataset, likely reflecting the effects of culture under homogeneous conditions.

**Figure 7.**
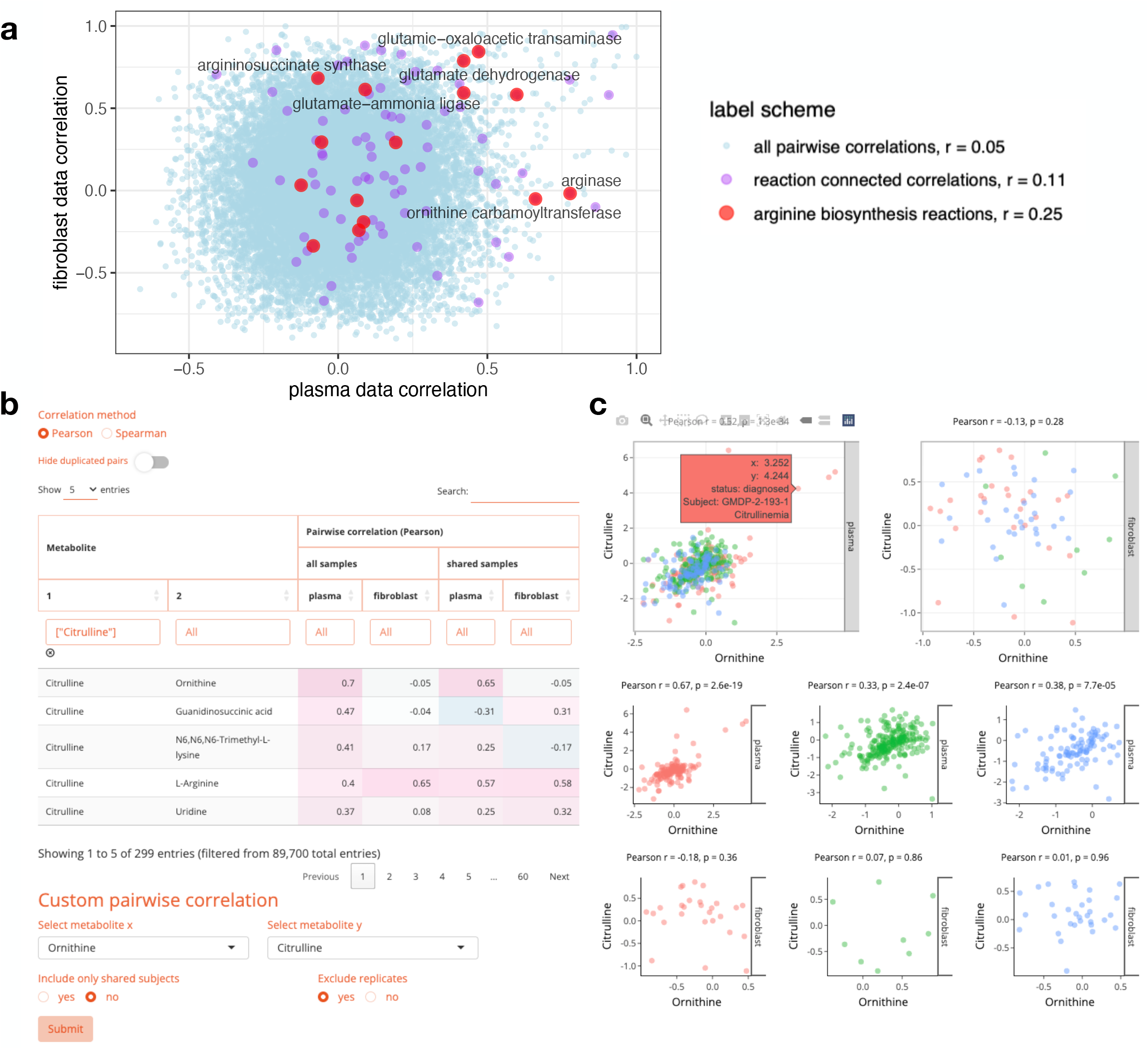
Metabolite pairwise correlations in plasma vs. fibroblast datasets. **a**, This scatter plot visualizes 44,850 pairwise correlation coefficients for 300 metabolites shared between fibroblast and plasma matrices. Each point represents a unique pair, with the correlation in fibroblast data plotted on the y-axis and the correlation in plasma data on the x-axis. Highlighted in purple are the correlations between metabolite pairs that are directly connected through a known reaction. More prominently featured in red are those pairs that participate specifically in the arginine biosynthesis pathway. Enzymes associated with large correlation coefficients within this pathway are labeled, to highlight reactions with strong inter-metabolite relationships. The plot reveals that reaction-connected correlations generally exhibit a positive correlation in both matrices. However, some reaction-connected pairs show strong correlation in one matrix but not the other, suggesting the differential presence or enzymatic activity within the arginine biosynthesis pathway between plasma and fibroblast environments. **b**, In the correlation table, ornithine is selected as the primary metabolite to identify its strongest correlations with other metabolites. Citrulline emerges with the highest correlation value, prompting further examination of their relationship. The table displays pairwise Pearson correlation coefficients for both plasma and fibroblast samples, with and without consideration of shared sample subsets. The ‘Hide duplicated pairs’ option is turned off to enable full visibility of all metabolite pairings. **c**, Custom pairwise correlation scatter plots comparing ornithine and citrulline levels across different subjects. The plots are stratified by diagnostic status, including diagnosed (red), unaffected (green), and unknown (blue).

The “Custom Metabolite Pairwise Correlations” feature of our webtool offers an interactive platform for comprehensive analysis of metabolite relationships in plasma and fibroblast datasets, providing researchers with the flexibility to explore correlations using data from either the subset of shared subjects or the entire cohort. The tool is especially adept at enabling users to discern the strength and directionality of correlations across these two distinct cellular environments. Within this analytical framework, Figure 7b showcases the correlation table, wherein ornithine is pre-selected to reveal its association with other metabolites. Citrulline is identified as having the most significant correlation with ornithine. This pairing reflects their well-established connection through the enzyme ornithine transcarbamylase (OTC), integral to the urea cycle.

Further illustrating the utility of the webtool, Figure 7c features custom pairwise correlation scatter plots that contrast ornithine and citrulline levels within various diagnostic groups differentiated by color, with diagnosed subjects in red, unaffected individuals in green, and those of unknown status in blue. Across all plasma-derived samples, a significant correlation is observed, yet this trend is notably absent in fibroblast samples, underscoring the lack of OTC expression in these cells. Within the diagnosed plasma subset, citrullinemia patients drive an even more pronounced correlation as a result of metabolic blockade by defects in the enzyme argininosuccinate synthetase (ASS), which intensifies the accumulation of both metabolites.

#### Pathway-Specific Metabolite Pairwise Correlations

Among the multitude of metabolite pairwise correlations, those encompassing metabolites within the same biochemical pathway often yield more coherent interpretations. The “Pathway-specific metabolite pairwise correlations” function enables users to explore correlations among metabolites within the same pathway. This feature provides options to filter for statistically significant correlations or those with direct biochemical linkages. Figure 8 focuses on the arginine biosynthesis pathway, highlighting correlations that exhibit differing patterns between plasma and fibroblast datasets. These variations may reflect the tissue-specific enzymatic activities, such as urea cycle enzymes active in the liver and gut but not in skin cells. This differential expression is exemplified by the notable correlation between ornithine and citrulline in plasma, absent in fibroblast data, aligning with the known tissue distribution of OTC.

**Figure 8.**
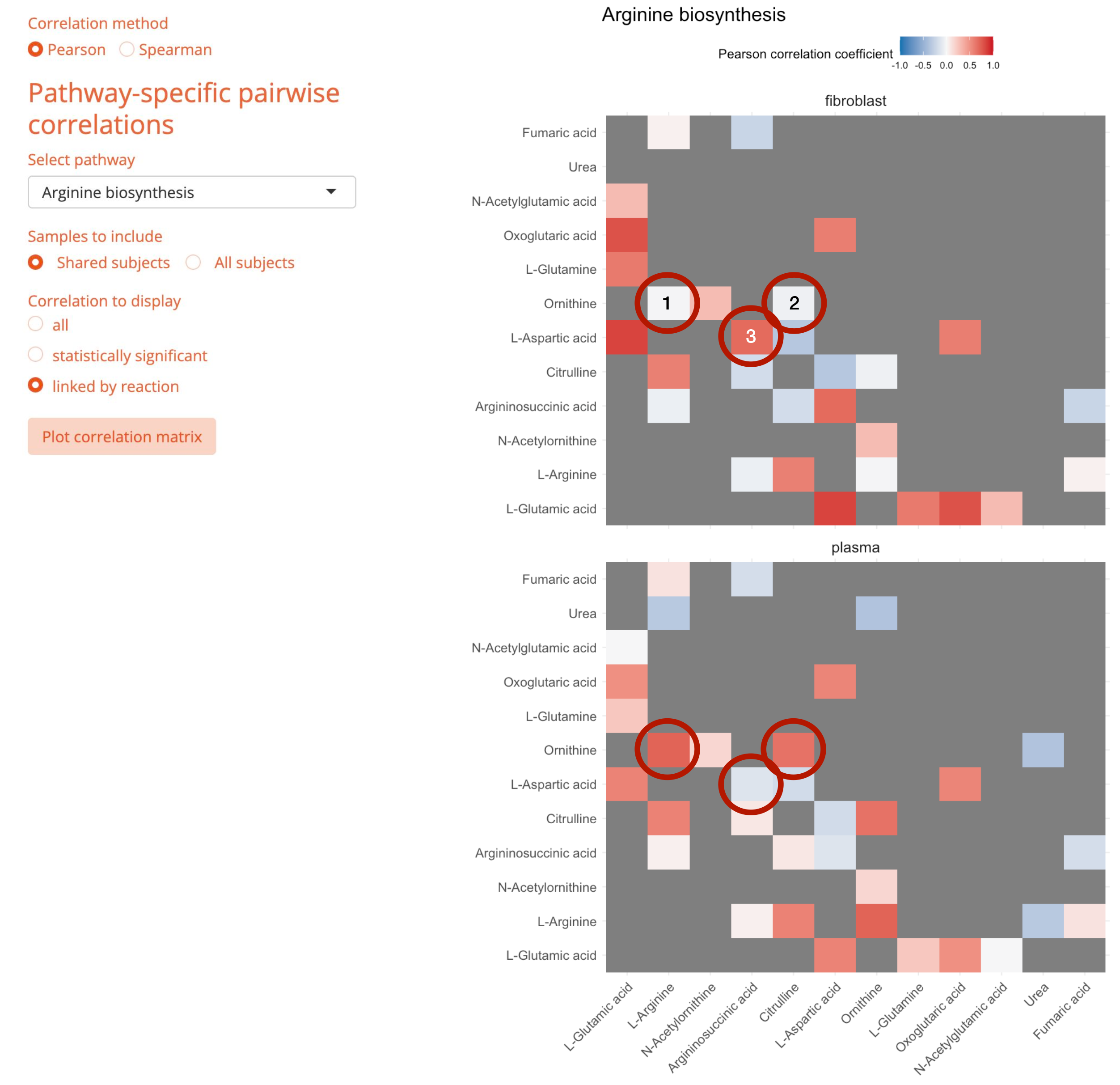
Metabolite pairwise correlation linked by reactions in the arginine biosynthesis pathways. The analysis outputs two heatmaps depicting the pairwise correlations of metabolites involved in the arginine biosynthesis pathway for plasma and fibroblast samples, respectively. The correlations are calculated using data from subjects shared between the two sample types and are specifically selected to show connections that are linked by known metabolic reactions within the pathway. The heatmaps are color-coded based on the Pearson correlation coefficient, ranging from -1 (perfect negative correlation, shown in blue) to +1 (perfect positive correlation, shown in red). Notable metabolic reactions with significant differences in correlation patterns between the plasma and fibroblast samples are encircled in red: (1) Arginase, which converts L-arginine to urea and ornithine, (2) Ornithine transcarbamylase, which converts ornithine to citrulline, and (3) Argininosuccinate synthase, which catalyzes the reaction between citrulline and aspartate to form argininosuccinate. These highlighted differences may reflect tissue-specific regulation or metabolic states in the sampled conditions. Note that only correlations linked by direct enzymatic reactions are displayed to focus on biologically relevant interactions within the pathway. This selective representation provides insights into the metabolic network’s complexity and its modulation in different biological matrices.

#### Reaction-Specific Metabolite Pairwise Correlations

Complementing the “Custom Metabolite Pairwise Correlations” and “Pathway-Specific Metabolite Pairwise Correlations,” the webtool “Reaction-Specific Metabolite Pairwise Correlations” enables users to interrogate the dynamic interplay between metabolites linked by a specific enzyme-catalyzed reaction. Users can navigate this tool by selecting a pathway or metabolite and refining their search down to the level of individual enzyme or gene names. Figure S10 illustrates the utility of this tool using the OTC reaction, previously discussed in Figure 7c. The interface not only displays the correlation in plasma and fibroblasts side by side, but also provides a schematic representation of the OTC reaction, indicating which metabolites are reactants or products, and the direction in which the reaction proceeds. This visual aid, complemented by KEGG pathway mappings, offers an intuitive understanding of the metabolic process, enhancing the researcher’s ability to interpret the data within the context of the broader metabolic network. It also allows the user to infer which reactions are present or absent in each matrix.

## Discussion and Conclusions

In this paper, we present a web tool tailored for exploration of plasma and fibroblast metabolomics data generated from a cohort enriched for patients with IEMs. While our companion study provides a detailed analysis of the data with wet lab validation[7], this paper focuses on the development and application of a user-friendly web application to explore the data. Our webtools are engineered to enable users to navigate and analyze the data with customizable features, providing insights into both systemic perturbations in plasma and cell-autonomous variability in fibroblast, in disease samples and healthy controls.

We have designed an array of user-guided functionalities, which include metabolic deviation analysis, metabolite clustering, and comparative studies between plasma and fibroblast metabolomes. These features are demonstrated through straightforward applications that extract insights from our data.

We acknowledge that our web application is specialized for our datasets and may not incorporate all possible analytical techniques. We also emphasize that because the data were gathered through a screening approach, individual metabolites should be validated by users on their own analytical systems prior to initiating extensive follow-on studies. Nonetheless, the structure of the webtool is intended to be versatile, potentially serving as a blueprint for other research groups to create similar tools adapted to their own datasets. Our hope is that by sharing our approach, we can contribute to a broader collective effort to decode the metabolic intricacies of diseases.

## Acknowledgements

We are grateful to the patients and families who provided samples for metabolomics analysis. R.J.D. is supported by the Howard Hughes Medical Institute Investigator Program, N.I.H. grant R35CA220449, the Baldridge Foundation, the Robert L. Moody, Sr. Faculty Scholar Award and the Eugene McDermott Distinguished Chair for the Study of Human Growth and Development. L.C. is supported by grants from NCI (R01CA285336) and CPRIT (RP250561). Metabolomics data were acquired in the Metabolomics Facility in Children’s Research Institute at UT Southwestern, which is supported by CPRIT grant RP180778. We wish to honor the late Mr. Gerardo Guevara for his pivotal role in sample biobanking that made this study possible.

