## Supplementary Figures S1-S10 for "An interactive web application for exploring human plasma and fibroblast metabolomics data from patients with inborn errors of metabolism"

**a**

Select matrix:
- (●) plasma
- ( ) fibroblast

Select dimension:
- (●) sample
- ( ) metabolite

Show 10 entries Search:

| ID | Subject | Dx | OMIM_ID | disease | centroid |
|---|---|---|---|---|---|
| All | All | All | All | All | All |
| GMDP-2-065 | GMDP-2-065-1 | 16p11.2 deletion syndrome | 611913 | CHROMOSOME 16p11.2 DELETION SYNDROME, 593-KB | yes |
| GMDP-2-074 | GMDP-2-074-1 | 22q11.2 deletion syndrome | 611867 | CHROMOSOME 22q11.2 DELETION SYNDROME, DISTAL | yes |
| GMDP-3-0136-1 | GMDP-3-0136-1 | ALPL mutation | 146300 | HYPOPHOSPHATASIA, ADULT | yes |
| GMDP-2-033_a | GMDP-2-033-1 | AMACR mutation | 614307 | ALPHA-METHYLACYL-CoA RACEMASE DEFICIENCY | yes |
| GMDP-2-138 | GMDP-2-138-1 | ASA | 207900 | ARGININOSUCCINIC ACIDURIA | yes |
| GMDP-3-0110-1 | GMDP-3-0110-1 | ASA | 207900 | ARGININOSUCCINIC ACIDURIA | yes |
| GMDP-2-089 | GMDP-2-089-1 | ASA | 207900 | ARGININOSUCCINIC ACIDURIA | yes |
| GMDP-3-0023-1 | GMDP-3-0023-1 | ASA | 207900 | ARGININOSUCCINIC ACIDURIA | yes |
| GMDP-2-038 | GMDP-2-038-1 | CUD | 212140 | CARNITINE DEFICIENCY, SYSTEMIC PRIMARY | yes |
| GMDP-2-037 | GMDP-2-037-1 | CUD | 212140 | CARNITINE DEFICIENCY, SYSTEMIC PRIMARY | yes |

Showing 1 to 10 of 697 entries Previous 1 2 3 4 5 ... 70 Next

Download

**b**

Select matrix:
- (●) plasma
- ( ) fibroblast

Select dimension:
- ( ) sample
- (●) metabolite

Show 15 entries Search:

| metabolite | Formula | Mass | HMDB | PubChem | KEGG |
|---|---|---|---|---|---|
| All | All | All | All | All | All |
| (2E,4E)-2,4-Hexadienoic acid | C6 H8 O2 | 112.0523 | HMDB0029581 | 643460 | |
| (±)-2-Hydroxy-4-(methylthio)butanoic acid | C5 H10 O3 S | 150.0346 | HMDB0037115 | 11427 | |
| 1-Aminocyclopropanecarboxylic acid | C4 H7 N O2 | 101.0475 | HMDB0036458 | 535 | C01234 |
| 1-Hydroxy-2-naphthoate | C11 H8 O3 | 188.0469 | | | C03203 |
| 1-Methyladenosine | C11 H15 N5 O4 | 281.1132 | HMDB0003331 | 27476 | C02494 |
| 1-Methylhistidine | C7 H11 N3 O2 | 169.0848 | HMDB0000001 | 92105 | C01152 |
| 10-Hydroxy-2-decenoic acid | C10 H18 O3 | 186.1254 | | | |
| 10-hydroxydecanoate | C10 H20 O3 | 188.1411 | | | |
| 10-undecenoate | C11 H20 O2 | 184.1461 | | | |
| 2',4'-Dihydroxyacetophenone | C8 H8 O3 | 152.0473 | HMDB0029659 | 6990 | C03663 |
| 2,6-Dihydroxypyridine | C5 H5 N O2 | 111.0323 | | | C03056 |
| 2-Aminophenol | C6 H7 N O | 109.0531 | | | C01987 |
| 2-Deoxyglucose | C6 H12 O5 | 164.0686 | HMDB0062477 | 108223 | |
| 2-Ethyl-2-Hydroxybutyric acid | C6 H12 O3 | 132.0787 | HMDB0001975 | 77199 | |
| 2-Hydroxybutyric acid | C4 H8 O3 | 104.0475 | HMDB0000008 | 11266 | C05984 |

Showing 1 to 15 of 412 entries Previous 1 2 3 4 5 ... 28 Next

Download

### Figure S1. Metadata page

**a**, Plasma data sample information. Links to to the corresponding Online Mendelian Inheritance in Man (OMIM) database entries are provided. The 'centroid' column indicates the representative sample for each subject, with 'yes' marking the samples selected as references in analyses where replicate exclusion is specified. **b**, Detailed metabolite information from the plasma dataset. Direct links to external databases for expanded data: the Human Metabolome Database (HMDB), PubChem, and the Kyoto Encyclopedia of Genes and Genomes (KEGG). This allows for easy cross-referencing and access to a wealth of chemical and biological information pertaining to each metabolite

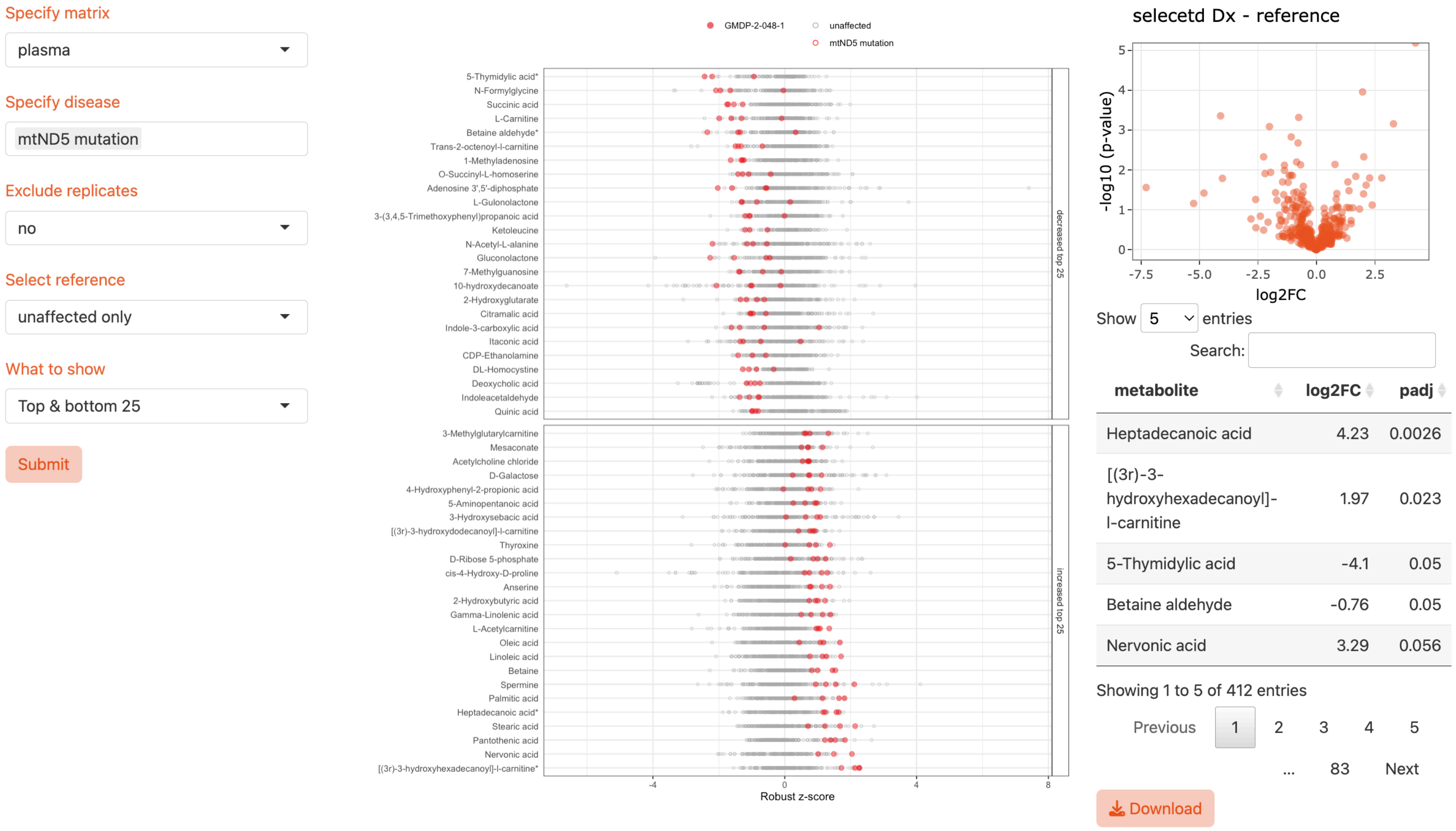


**Figure S2. Additional usage examples of Dx vs. reference comparison**
Comparison between mtND5 mutation and the unaffected subjects from the fibroblast data, showing the most accumulated and depleted metabolites. Metabolites are sorted based on the magnitude of deviation from the reference group, providing a ranking of metabolic disturbances associated with each condition.

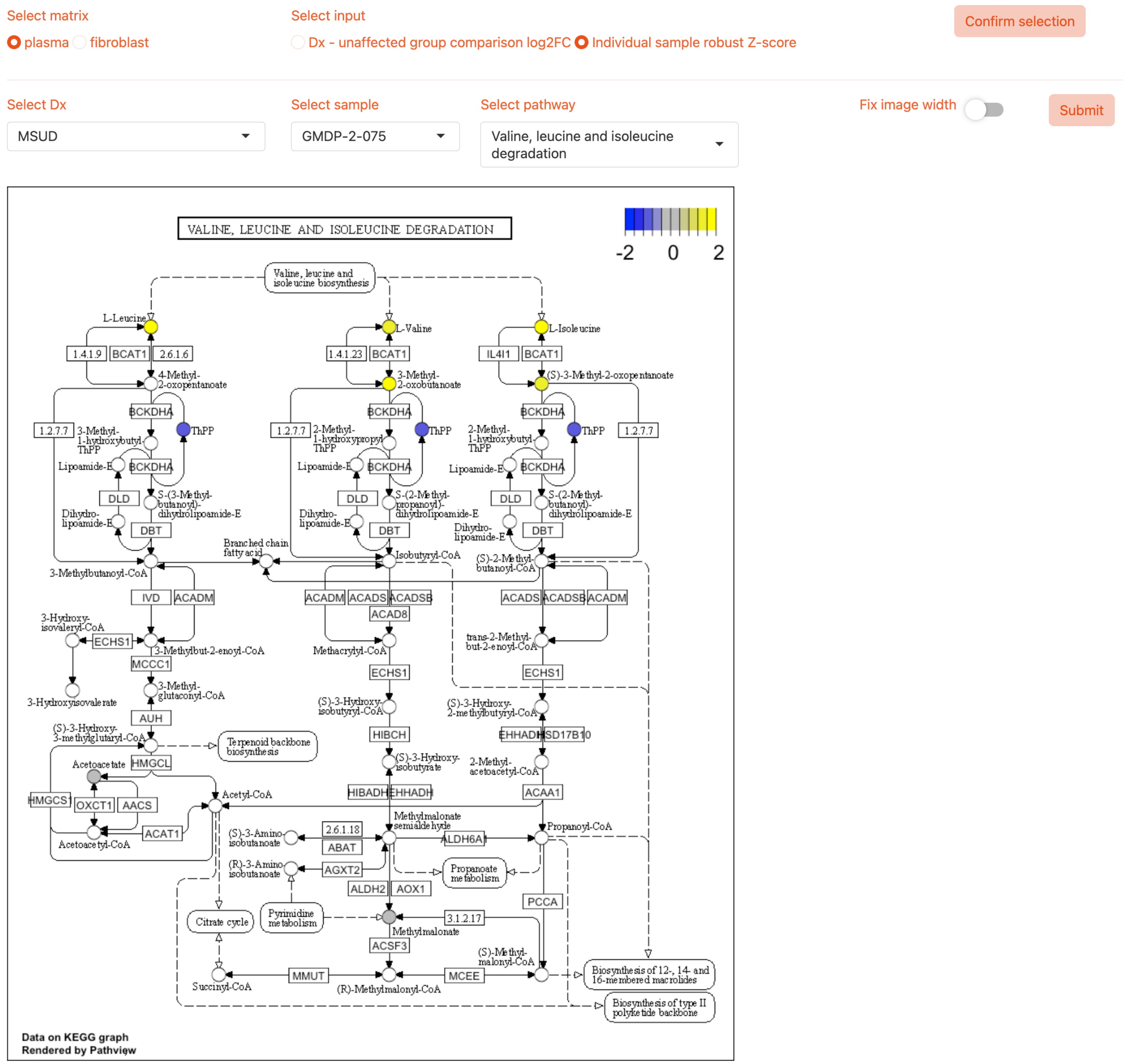


**Figure S3. Robust Z-score from an MSUD subject projected onto the BCAA catabolic pathway**

This visualization maps the robust z-scores of metabolites onto the branched-chain amino acid (BCAA) catabolism pathway for a subject with Maple Syrup Urine Disease (MSUD), identified as GMDP-2-075. The pathway diagram highlights the biochemical steps involved in the metabolism of BCAAs — valine, leucine, and isoleucine — whose breakdown is impaired in MSUD. Metabolites assessed in the study are marked within the pathway, with color intensity reflecting the magnitude of the robust z-score, indicating their relative concentration in the MSUD subject's sample.

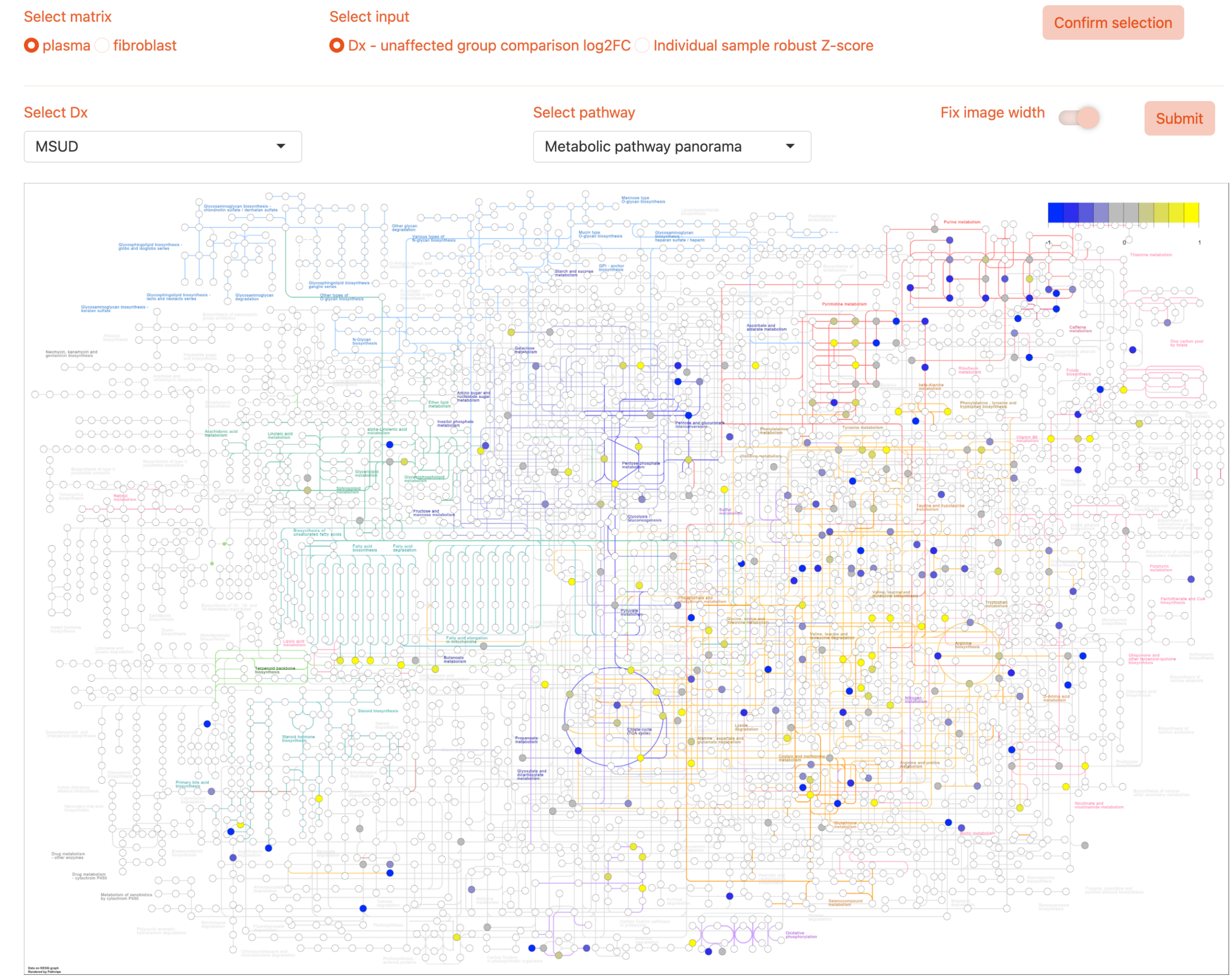


**Figure S4. Metabolic pathway panorama view for MSUD - reference log2FC**
This panoramic view of the metabolic pathways illustrates the differences in metabolite concentrations between Maple Syrup Urine Disease (MSUD) subjects and healthy controls. The input for this plot is the log2 fold change (log2FC), calculated using a linear mixed-effects model to compare the metabolic profiles. The 'Fix image width' feature has been enabled to fit the entire pathway panorama within the display frame; disabling this feature allows for a zoomed-in view for more detailed examination.

**a**

Select matrix

fibroblast

Exclude replicates

yes

Exclude samples with Dx

unaffected

unknown

Submit

Show 15 entries

Search:

| ID | Dx | metabolite | z-score |
|---|---|---|---|
| All | All | All | All |
| GMDP-4-0031-4_a | GA1 | Glutaric acid | 11.48 |
| GMDP-4-0031-1_a | GA1 | Glutaric acid | 10.08 |
| MEPAN-1_b | MECR mutation | 4-acetylbutyrate | 9.33 |
| VIG-LIPT1_c | LIPT1 mutation | 4-acetylbutyrate | 9.01 |
| BCH-1089_b | ODC1 mutation | 4-acetylbutyrate | 7.87 |
| GMDP-3-0041-1_b | EARS2 mutation | 4-acetylbutyrate | 7.61 |
| 1095_c | mtTL1 mutation | 4-acetylbutyrate | 6.86 |
| LIPT2-sib-2_a | LIPT2 mutation | 4-acetylbutyrate | 6.65 |
| LIPT2-sib-2_a | LIPT2 mutation | Mevalonic acid | 6.33 |
| MEPAN-1_b | MECR mutation | Hydroxyisocaproic acid | 6.03 |
| CHOP-F66S_a | PCK1 mutation | 4-acetylbutyrate | 5.95 |
| BCH-1089_b | ODC1 mutation | Hydroxyisocaproic acid | 5.75 |
| LIPT2-sib-2_a | LIPT2 mutation | Hydroxyisocaproic acid | 5.65 |
| VIG-LIPT1_c | LIPT1 mutation | Hydroxyisocaproic acid | 5.5 |
| 1095_c | mtTL1 mutation | Hydroxyisocaproic acid | 5.35 |

Showing 1 to 15 of 139 entries

Previous 1 2 3 4 5 ... 10 Next

**b**

Select matrix

plasma

Exclude replicates

yes

Exclude samples with Dx

unaffected

unknown

Submit

Show 15 entries

Search:

| ID | Dx | metabolite | z-score |
|---|---|---|---|
| All | All | ["Methylmalonic acid"] | All |
| GMDP-2-131 | MMA | Methylmalonic acid | 7.27 |
| GMDP-2-052 | MMA | Methylmalonic acid | 6.84 |
| GMDP-2-053_a | MMA | Methylmalonic acid | 6.79 |
| GMDP-2-131-4 | MMA | Methylmalonic acid | 5.82 |
| GMDP-2-021_a | CblC | Methylmalonic acid | 4.69 |
| GMDP-2-173 | CblC | Methylmalonic acid | 4.52 |
| GMDP-2-128 | CblC | Methylmalonic acid | 3.55 |
| GMDP-2-018_a | MMA | Methylmalonic acid | 3.46 |
| GMDP-4-0011-1 | CblC | Methylmalonic acid | 3.27 |
| GMDP-4-0008-1 | CblC | Methylmalonic acid | 3.11 |
| GMDP-2-130_b | CblC | Methylmalonic acid | 3.09 |
| GMDP-3-0149-2 | unaffected | Methylmalonic acid | -3.01 |
| GMDP-2-190 | GA1 | Methylmalonic acid | -3.27 |
| GMDP-3-0121-1 | unknown | Methylmalonic acid | -3.99 |

Showing 1 to 14 of 14 entries (filtered from 1,039 total entries)

Previous 1 Next

**Figure S5. Metabolic outliers in the GMDP datasets**

**a**, Metabolic outliers in the fibroblast dataset in descending order of robust z-scores. Notably, the metabolite 4-acetylbutyrate appears recurrently among various diagnosed samples, including those with mitochondrial defects. While samples from unaffected and unknown diagnoses were included in the dataset, the top outliers are exclusively from diagnosed subjects. **b**, Top z-score deviations for methylmalonic acid in the plasma dataset. The list reveals that all positive outliers with elevated levels of this metabolite are from subjects diagnosed with Methylmalonic Acidemia (MMA) or Cobalamin C Deficiency (CblC).

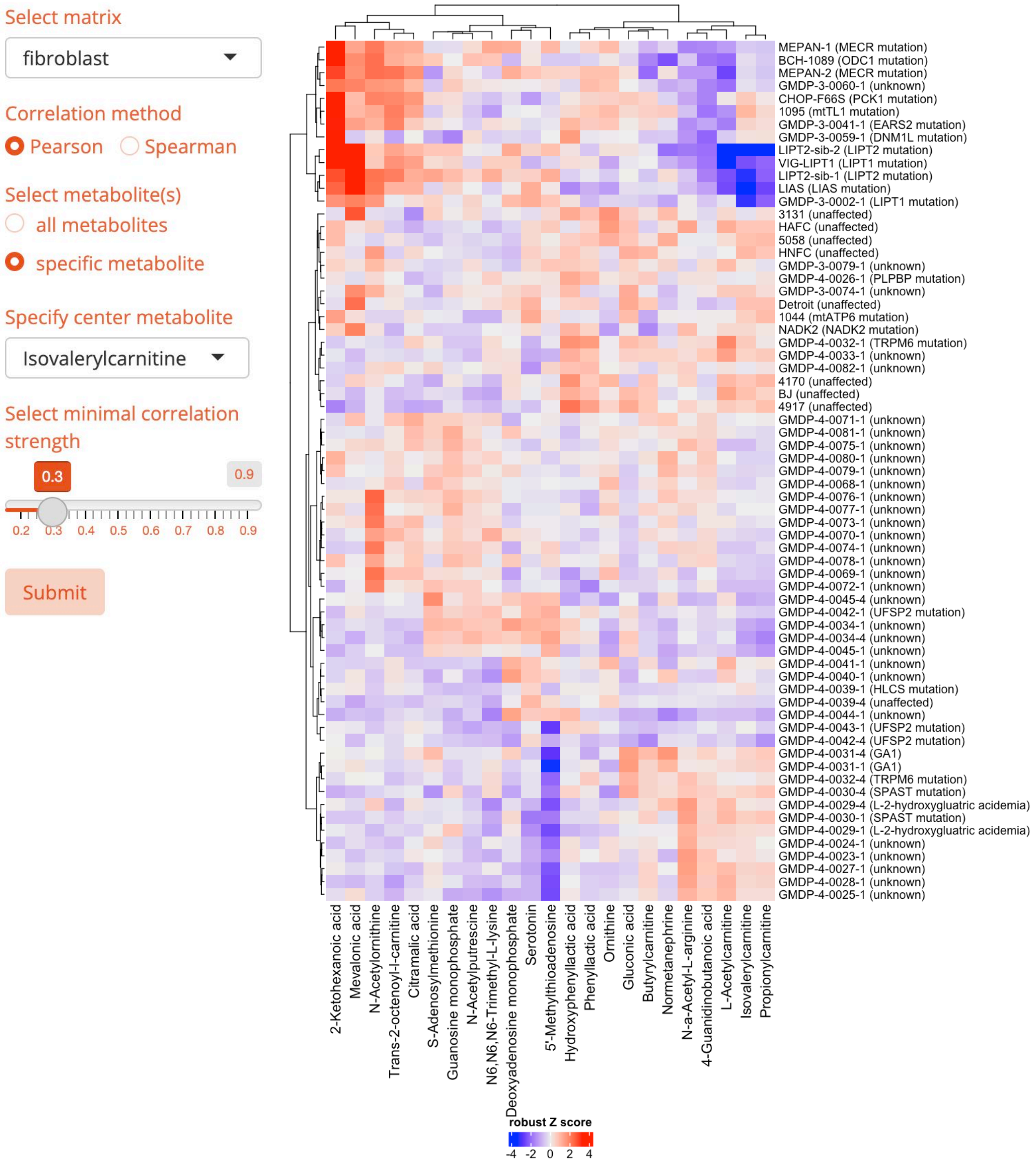


**Figure S6. Fibroblast metabolites that correlate with isovalerylcarnitine**

This heatmap visualizes the correlation of isovalerylcarnitine with 22 related metabolites in fibroblast samples, chosen for their absolute correlation strength above 0.3. The robust z-scores for these metabolites form the basis of the clustered heatmap, highlighting the relationships within centroid samples excluding replicates. Each subject's diagnosis is indicated alongside their ID. Notably, isovalerylcarnitine shows a strong association with other short-chain acylcarnitines.

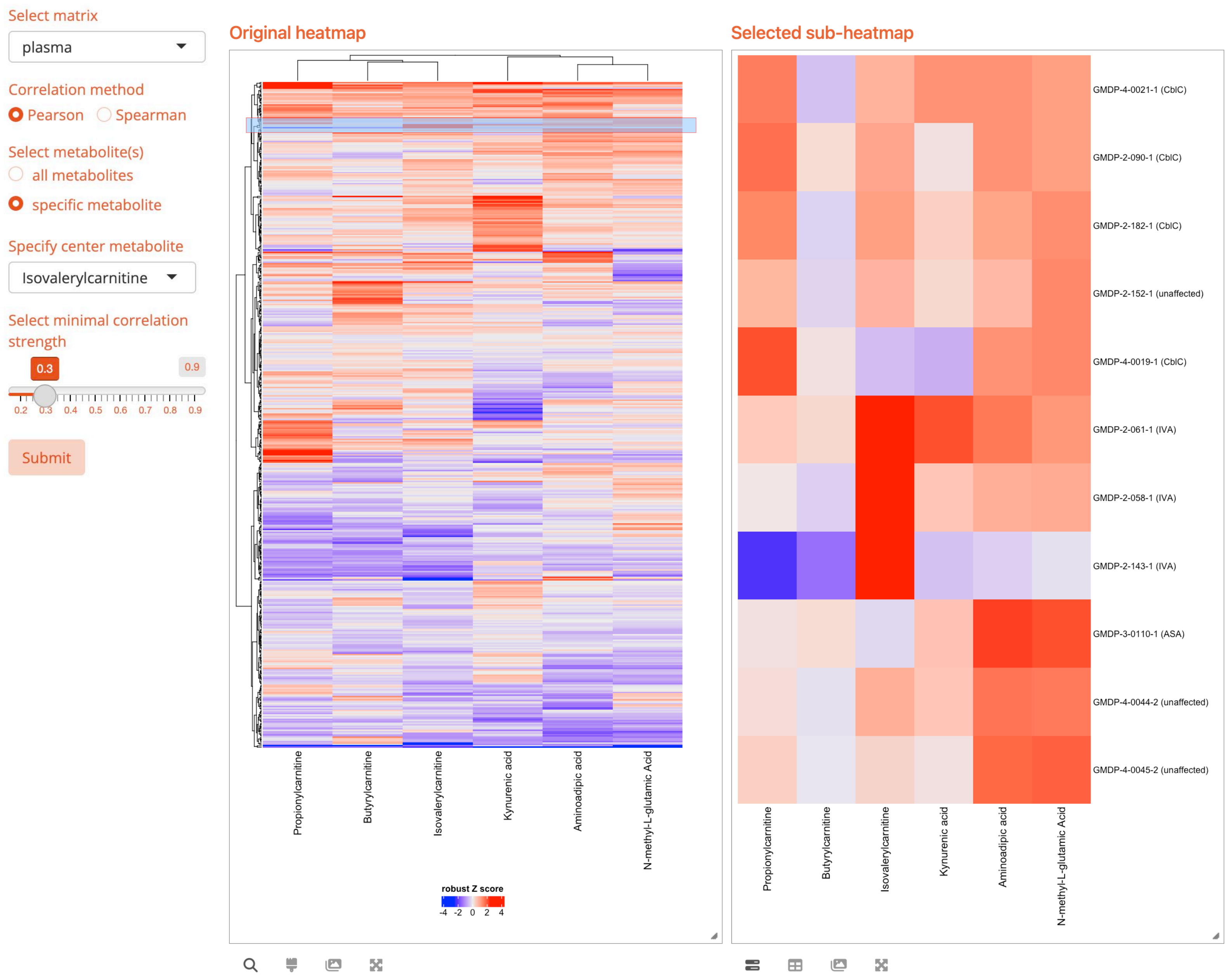


**Figure S7. Plasma metabolites that correlate with isovalerylcarnitine**

A heatmap illustrates the correlation of isovalerylcarnitine with five metabolites in plasma, selected for correlation strength above 0.3. The robust z-scores inform the clustering within centroid samples, excluding replicates. A magnified sub-heatmap feature allows for detailed examination of specific regions. Isovalerylcarnitine is notably clustered with other short-chain acylcarnitines, suggesting a shared metabolic relationship.

**a**

Specify matrix

plasma

Specify dimension

metabolite

Exclude replicates

yes

Samples to include

- [x] diagnosed
- [x] unknown
- [x] unaffected

Choose ranking distance

- ○ Euclidean distance
- ● Absolute correlation distance

Correlation method

● Pearson ○ Spearman

Submit

Show 10 entries

Search:

| metabolite | metabolite_neighbor | distance | r | pv | n |
|---|---|---|---|---|---|
| ["Propionylcarr | All | All | All | All | All |
| Propionylcarnitine | Methylmalonic acid | 0.55 | 0.45 | 0 | 474 |
| Propionylcarnitine | Isovalerylcarnitine | 0.58 | 0.42 | 0 | 474 |
| Propionylcarnitine | Butyrylcarnitine | 0.61 | 0.39 | 0 | 474 |
| Propionylcarnitine | Aminoadipic acid | 0.71 | 0.29 | 1.3e-10 | 474 |
| Propionylcarnitine | N-Acetyl-L-alanine | 0.74 | 0.26 | 2e-8 | 468 |
| Propionylcarnitine | L-Acetylcarnitine | 0.75 | 0.25 | 3.3e-8 | 474 |
| Propionylcarnitine | L-Homocysteine thiolactone | 0.77 | 0.23 | 2.3e-7 | 474 |
| Propionylcarnitine | 5-Hydroxy-L-tryptophan | 0.78 | 0.22 | 0.0000012 | 474 |
| Propionylcarnitine | L-Kynurenine | 0.79 | 0.21 | 0.0000031 | 474 |
| Propionylcarnitine | (±)-2-Hydroxy-4-(methylthio)butanoic acid | 0.81 | 0.19 | 0.000026 | 474 |

Showing 1 to 10 of 411 entries (filtered from 169,332 total entries)

Previous 1 2 3 4 5 … 42 Next

**b**

Specify matrix

fibroblast

Specify dimension

sample

Exclude replicates

yes

Samples to include

- [x] diagnosed
- [x] unknown
- [x] unaffected

Choose ranking distance

- ● Euclidean distance
- ○ Absolute correlation distance

Submit

Show 15 entries

Search:

| sample_ID | sample_Dx | neighbor_ID | neighbor_Dx | distance |
|---|---|---|---|---|
| VIG-LIPT1 | All | All | All | All |
| VIG-LIPT1_c | LIPT1 mutation | LIPT2-sib-2_a | LIPT2 mutation | 13.52 |
| VIG-LIPT1_c | LIPT1 mutation | 1095_c | mtTL1 mutation | 13.9 |
| VIG-LIPT1_c | LIPT1 mutation | GMDP-3-0041-1_b | EARS2 mutation | 13.93 |
| VIG-LIPT1_c | LIPT1 mutation | CHOP-F66S_a | PCK1 mutation | 15.91 |
| VIG-LIPT1_c | LIPT1 mutation | LIPT2-sib-1_a | LIPT2 mutation | 16.56 |
| VIG-LIPT1_c | LIPT1 mutation | BCH-1089_b | ODC1 mutation | 16.57 |
| VIG-LIPT1_c | LIPT1 mutation | GMDP-3-0060-1_b | unknown | 17.07 |
| VIG-LIPT1_c | LIPT1 mutation | MEPAN-1_b | MECR mutation | 17.99 |
| VIG-LIPT1_c | LIPT1 mutation | LIAS_b | LIAS mutation | 18.66 |
| VIG-LIPT1_c | LIPT1 mutation | MEPAN-2_c | MECR mutation | 19.02 |
| VIG-LIPT1_c | LIPT1 mutation | GMDP-3-0002-1_f | LIPT1 mutation | 20.36 |
| VIG-LIPT1_c | LIPT1 mutation | 1044_b | mtATP6 mutation | 20.45 |
| VIG-LIPT1_c | LIPT1 mutation | HNFC_c | unaffected | 20.83 |
| VIG-LIPT1_c | LIPT1 mutation | 5058_a | unaffected | 21.35 |
| VIG-LIPT1_c | LIPT1 mutation | GMDP-3-0079-1_b | unknown | 21.68 |

Showing 1 to 15 of 66 entries (filtered from 4,422 total entries)

Previous 1 2 3 4 5 Next

**Figure S8. Investigating metabolite and sample similarity with “Neighbor distance table”**

**a**, Plasma metabolites closely related to propionylcarnitine, ranked by absolute correlation distance. This analysis identifies metabolites that exhibit similar patterns of variation. **b**, Fibroblast samples with profiles similar to the sample VIG-LIPT1_c, ordered by Euclidean distance. This comparison highlights samples with closely matching metabolic profiles.

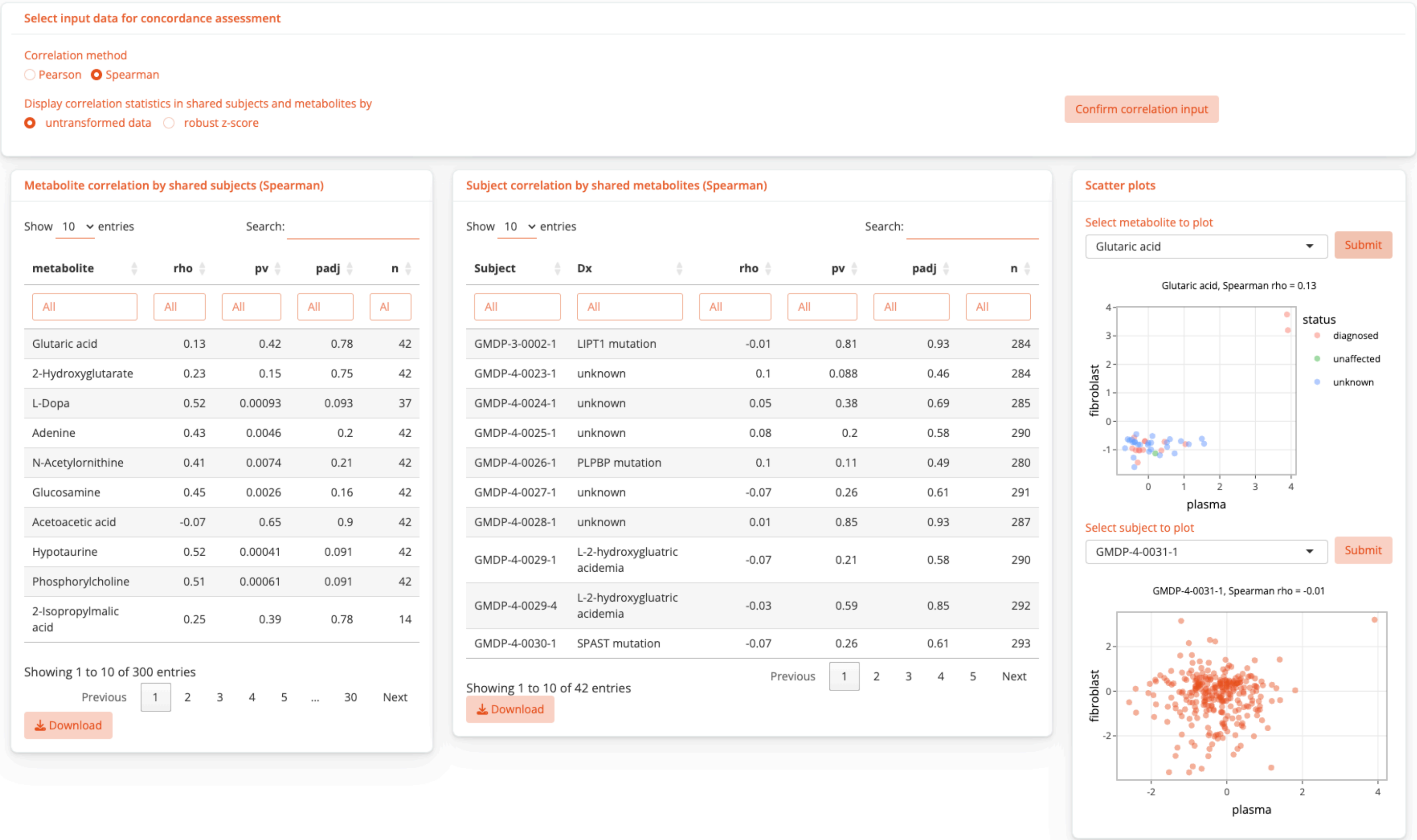


| metabolite | rho | pv | padj | n |
|---|---|---|---|---|
| All | All | All | All | Al |
| Glutaric acid | 0.13 | 0.42 | 0.78 | 42 |
| 2-Hydroxyglutarate | 0.23 | 0.15 | 0.75 | 42 |
| L-Dopa | 0.52 | 0.00093 | 0.093 | 37 |
| Adenine | 0.43 | 0.0046 | 0.2 | 42 |
| N-Acetylornithine | 0.41 | 0.0074 | 0.21 | 42 |
| Glucosamine | 0.45 | 0.0026 | 0.16 | 42 |
| Acetoacetic acid | -0.07 | 0.65 | 0.9 | 42 |
| Hypotaurine | 0.52 | 0.00041 | 0.091 | 42 |
| Phosphorylcholine | 0.51 | 0.00061 | 0.091 | 42 |
| 2-Isopropylmalic acid | 0.25 | 0.39 | 0.78 | 14 |

| Subject | Dx | rho | pv | padj | n |
|---|---|---|---|---|---|
| All | All | All | All | All | All |
| GMDP-3-0002-1 | LIPT1 mutation | -0.01 | 0.81 | 0.93 | 284 |
| GMDP-4-0023-1 | unknown | 0.1 | 0.088 | 0.46 | 284 |
| GMDP-4-0024-1 | unknown | 0.05 | 0.38 | 0.69 | 285 |
| GMDP-4-0025-1 | unknown | 0.08 | 0.2 | 0.58 | 290 |
| GMDP-4-0026-1 | PLPBP mutation | 0.1 | 0.11 | 0.49 | 280 |
| GMDP-4-0027-1 | unknown | -0.07 | 0.26 | 0.61 | 291 |
| GMDP-4-0028-1 | unknown | 0.01 | 0.85 | 0.93 | 287 |
| GMDP-4-0029-1 | L-2-hydroxygluatric acidemia | -0.07 | 0.21 | 0.58 | 290 |
| GMDP-4-0029-4 | L-2-hydroxygluatric acidemia | -0.03 | 0.59 | 0.85 | 292 |
| GMDP-4-0030-1 | SPAST mutation | -0.07 | 0.26 | 0.61 | 293 |

**Figure S9. Metabolomic concordance in plasma and fibroblasts using with samples from shared subjects**

Left: These tables display metabolic correlations within shared subjects and between shared metabolites, available for review and download. Nominal p-values are displayed. Right: scatter plot illustrating selected examples of metabolic correlations between fibroblast and plasma profiles, emphasizing variations attributable to specific disease states or metabolites. For instance, the plot for glutaric acid shows a correlation primarily influenced by samples from patients diagnosed with Glutaric Aciduria Type 1.

Correction method

Correlation method

◉ Pearson ○ Spearman

### Reaction-specific pathway correlation(s)

Samples to include

○ Shared subjects ◉ All subjects

Select reaction by

◉ pathway ○ compound

Select pathway

Arginine biosynthesis

Specify reaction by

○ Enzyme name ◉ Gene name

Select reaction

OTC (R01398)

Carbamoyl-phosphate:L-ornithine carbamoyltransferase
Carbamoyl phosphate + L-Ornithine <=> Orthophosphate + L-Citrulline
C00169 + C00077 <=> C00009 + C00327

Submit

| KEGG | metabolite | plasma | fibroblast |
|---|---|---|---|
| C00077 | L-Ornithine | true | true |
| C00327 | L-Citrulline | true | true |
| C00009 | Orthophosphate | false | false |
| C00169 | Carbamoyl phosphate | false | false |

Availability of reaction-related metabolites

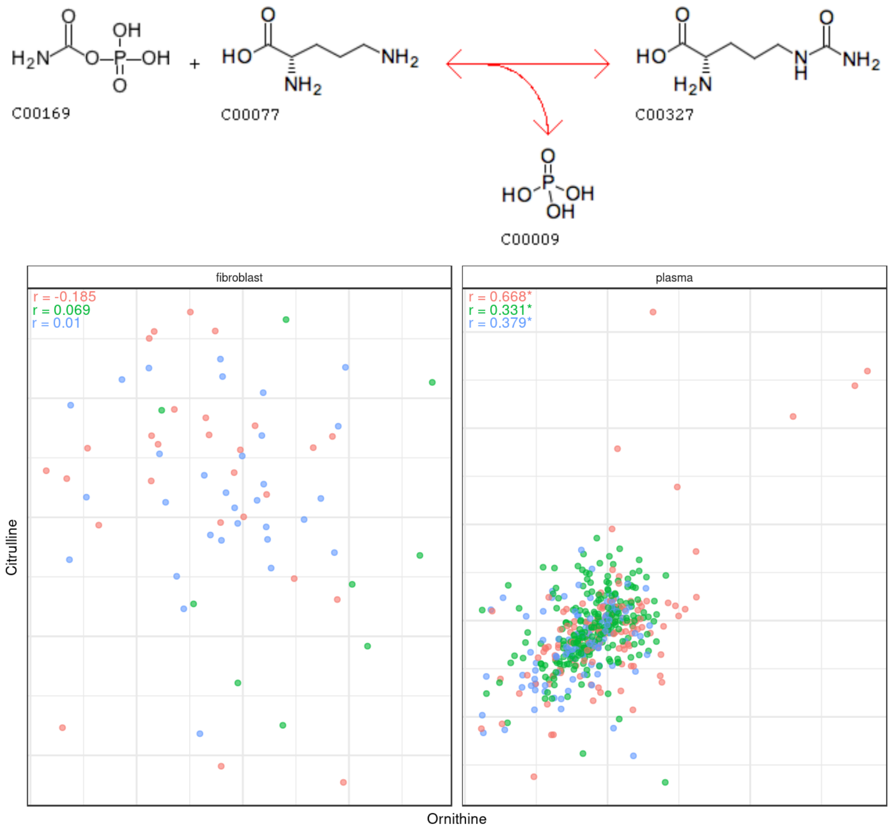


**Figure S10. Examination of metabolite correlation within specific metabolic reactions**
This interface allows users to analyze correlations based on selected metabolic reactions from specified pathways or individual compounds. The system limits selection to those reactions and compounds available on the metabolomics platform. Upon choosing "All subjects," the platform generates correlation statistics that differentiate diagnosis statuses: red dots represent diagnosed subjects, green for unaffected, and blue for unknown diagnoses. The example shown illustrates the correlation of compounds involved in the ornithine transcarbamylase (OTC) reaction within the arginine biosynthesis pathway, with separate correlation scatter plots for fibroblast and plasma samples. Note that compounds not profiled by the platform are excluded, ensuring focused analysis on data-supported interactions.
